# Fungal organic acid uptake of mineral derived K is dependent on distance from carbon hotspot

**DOI:** 10.1101/2023.03.17.533228

**Authors:** Arunima Bhattacharjee, Dusan Velickovic, Jocelyn A. Richardson, Sneha P. Couvillion, Gregory W. Vandergrift, Odeta Qafoku, Michael J. Taylor, Janet K. Jansson, Kirsten S. Hofmockel, Christopher R. Anderton

**Affiliations:** Environmental Molecular Sciences Division, Earth and Biological Sciences Directorate, Pacific Northwest National Laboratory, Richland, WA 99352; Stanford Synchrotron Radiation Lightsource, SLAC National Accelerator Laboratory, Menlo Park, CA 94025; Biological Sciences Division, Earth and Biological Sciences Directorate, Pacific Northwest National Laboratory, Richland, WA 99352

## Abstract

Fungal mineral weathering processes regulate the bioavailability of inorganic nutrients from mineral surfaces to organic matter and increase the bioavailable fraction of nutrients. Fungal mineral weathering strategies can be classified as biomechanical (direct) or biochemical (indirect). In the case of fungal uptake of mineral nutrients through indirect weathering, it is widely hypothesized that uptake of mineral derived nutrients occurs through organic acid chelation, but such processes have not been directly visualized. This is in part due to challenges in probing the complex and heterogeneous soil environment. Here, using an epoxy-based mineral doped soil micromodel platform that we developed, which mimics soil mineralogy and structure, it permitted us to visualize and spatially probe the molecular mechanisms of mineral weathering. Mass spectrometry imaging revealed differences in the distribution of fungal exudates, citric acid and tartaric acid, on the soil micromodels in presence of minerals. Citric acid was detected closer to the nutrient rich inoculation point, whereas tartaric acid was highly abundant away from inoculation point. This suggested that the organic acid exuded by the fungi depended on the proximity from the carbon rich organic substrate at the point of inoculation. Using a combination of X-ray fluorescence and X-ray near edge structure analysis, we identified citric acid- and tartaric acid-bound K within fungal hyphae networks grown in the presence of minerals. Combined our results provide direct evidence that fungi uptake and transport mineral derived nutrient organic acid chelation. The results of this study provided unprecedented visualization of fungal weathering of soil minerals and hyphal K^+^ transport, while resolving the indirect weathering mechanism of fungal K uptake from mineral interfaces.

## Introduction

Soil organic carbon distribution is spatially heterogeneous, occurring in discontinuous patchy islands or carbon hotspots in soil that are accessed by microbial communities and plants for growth and development(1–5). The discontinuous carbon hotspots create differences in microbial community phenotype and exudation of metabolites required for uptake of nutrients or intra and interspecies interactions(6). For example, a greater access to nutrients including carbon, induces increase in microbial population, while decreasing microbial interactions for resource access. On the other hand, there is increased interactions and competition for resources in a carbon limited environment(7). Extensive research has been conducted on microbial phenotype changes as a function of varying nutrient availability in liquid and bulk soil cultures(8, 9). However, how microbes spatially navigate the patchy network of carbon hotspots, while interfacing with soil minerals and extracting nutrients for continued community development in between carbon hotspots has not been extensively studied.

Soil minerals form a huge reservoir of inorganic nutrients for microbial communities. Soil microorganisms access mineral derive inorganic nutrients through biomechanical (direct) or biochemical (indirect) weathering of minerals(10). Most microbially induced biochemical means of mineral weathering involve production of low molecular weight polycarboxylic organic acids such as citric, malic, oxalic, and maleic acids, as well as extracellular polymeric substances(11, 12). These organic acids create microbially accessible pools of bioavailable inorganic nutrients such as potassium, calcium, iron etc. For example, most potassium (K), a macronutrient within primary minerals is not available for microbial uptake, but microbial weathering can create accessible K pools for microbial uptake. Investigating microbial mineral weathering spatially in relation to carbon hotspots is essential in elucidating mechanisms of biotic driven cycling of mineral derived inorganic nutrients and microbially derived carbon retention on weathered mineral surfaces. However, spatial investigation of mechanisms of inorganic nutrient uptake and cycling in soil is complicated by the heterogeneous nature of the soil landscape, among other limitations.

Previously, we demonstrated a mineral doped soil micromodel platform for studying fungal bridging of carbon rich hotspots as well as fungal derived mineral weathering(13). This system reduces the complexity of soil while incorporating sentinel elements of soil necessary for testing specific hypothesis. In this work, we used K – feldspar mineral doped soil micromodels and a saprotrophic fungus, *Fusarium sp. DS 682*(14) to demonstrate that fungal organic acid production changes as a function of distance from a carbon hotspot. Consequently, the organic acid uptake of inorganic nutrients from minerals changes due to different types of organic acids being exuded by fungal hyphae. Specifically, we show that the uptake of mineral derived K from mineral interfaces into fungal hyphae occurs through citric and tartaric acid chelation and the uptake organic acid type changes as a function of distance from a carbon rich hotspot.

## Results

### Organic acid production by *Fusarium sp. DS 682* in response to the presence of minerals

Organic acid production in *Fusarium* in relation to mineral weathering has not been thoroughly studied. Here, we analyzed organic acids produced by *Fusarium sp. DS 682* during growth on agar plates with and without minerals(13). We observed that this saprotrophic soil fungi produces several organic acids such as citric acid, fumaric acid, malic acid, oxalic acid, and succinic acid (Fig. 1A). The relative concentrations of organic acids were higher in the treatment group with minerals in comparison to those grown in the absence of minerals.

**Fig. 1.**
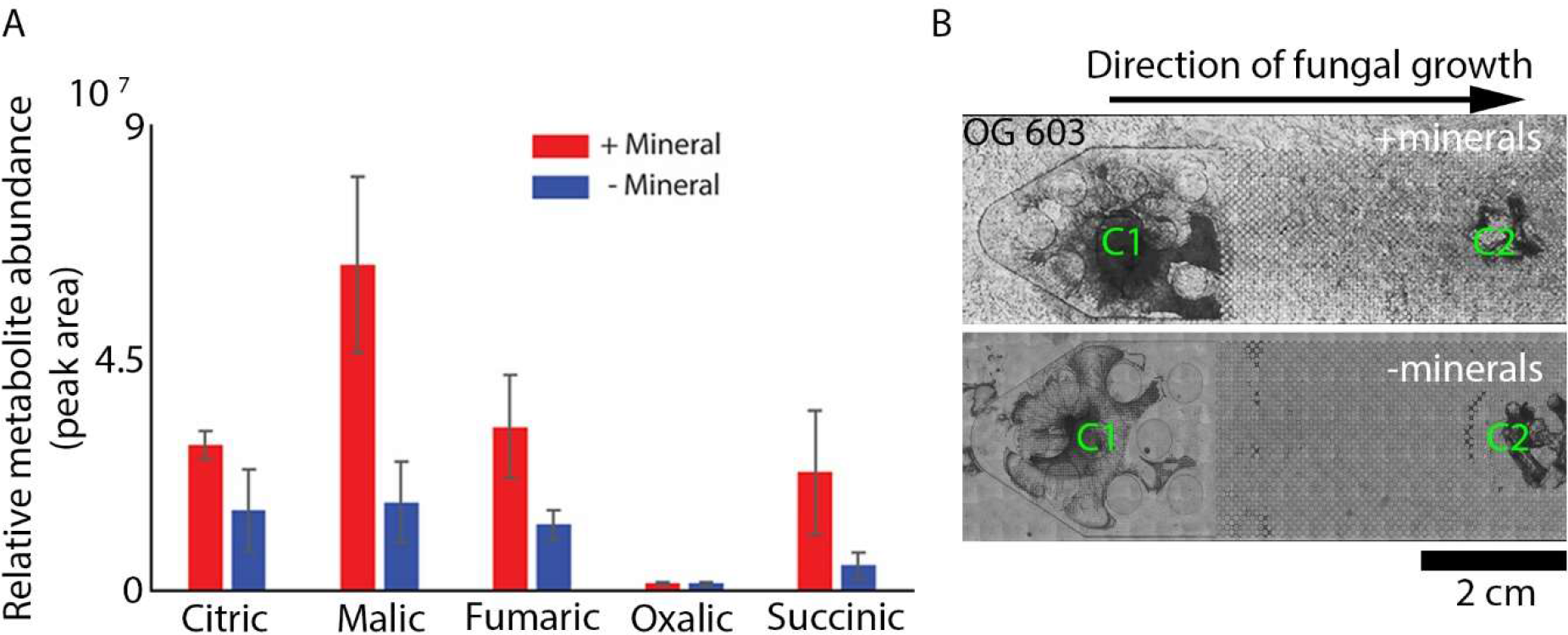
Fungal organic acid production increases in the presence of minerals. A) GCMS data of fungal biomass grown in presence (+Mineral) and absence (-Mineral) of minerals on potato dextrose agar surfaces shows an increase in relative concentration of organic acid in presence of minerals. B) Fungal growth occurs over 7 days on OG 603 polymer micromodels in presence of minerals (+minerals) between nutrient rich PDA plugs at two ends of the micromodel (C1 and C2) (top image). Fungal hyphae growth on micromodels without minerals (-minerals) (bottom image).

Fungal growth on agar surfaces demonstrated an increase in the relative concentrations of organic acids in the presence of minerals, indicating the potential these organic acids play in acquisition of mineral derived nutrients. However, agar plate cultures are not an accurate representation of a soil microenvironment, which comprise of soil aggregates, varying porosities, and different mineralogy. In our previous work, we created soil micromodels that mimic soil porosity and mineralogy, in effort to demonstrate fungal induced mineral weathering, extraction, and transport of potassium (K)(13). Here, we created mineral doped micromodels for detection of fungal organic acids by adding solid phase kaolinite mineral into a biocompatible and UV curable polymer OG603, similar to our recent work(15), before molding into the shape of the micromodel channel. The minerals embedded into the polymer were made accessible on the micromodel channel surface through a O2 plasma dry etching process (Fig. 1B and S1). The kaolinite mineral was previously used to study fungal growth and thigmotropism in mineral doped micromodels, and it is enriched with 7% K-feldspar and 4% mica(13). Fungi was inoculated in one end of the micromodel channel along with two organic nutrient rich plugs composed of potato dextrose agar (PDA) (C1 and C2, Fig. 1B) with the micromodel pore space in between. The micromodel channel space was kept unsaturated and the only nutrient sources between the PDA plugs were inorganic mineral-bound nutrients. We observed increased fungal growth within the micromodel channels with minerals in 7 days and 30 days compared to micromodel channels with no minerals, which is consistent with our previous results (Fig.1B)(13).

### Secreted fungal organic acid distribution is spatially heterogeneous

The spatial distribution of organic acid produced by *Fusarium sp. DS 682* was analyzed using matrix-assisted laser desorption/ionization mass spectrometry imaging (MALDI-MSI) on the OG603 micromodel surfaces after fungal growth. Apart from citric acid, we also detected that fumaric, malic, tartaric, oxoadipic, and galactonic/gluconic acid (undisguisable isomers) was produced during fungal growth on mineral doped micromodel surfaces (Figs. 2 and S2). Citric, malic and fumaric acids were concentrated closer to the C-rich inoculation point, while tartaric, oxoadipic and galactonic/gluconic acids increased in abundance with distance from the C hotspot. In comparison, the mineral free control contained relatively lower concentrations of fumaric, malic, and citric acid, and the localization of these organic acids remained at and around the inoculation point (Fig. S3). The differences in distribution of fungal organic acids between micromodels with and without minerals suggests that a change in the type of organic acid production occurs in response to the presence of minerals. These differences were not observable through our bulk analysis of fungal biomass from agar plates (Fig. 1), which provides no spatial context to the distribution of these molecules. Moreover, we observed differences in the spatial distribution of organic acids on the mineral doped micromodels alone. Tartaric acid, oxoadipic acid, and galactonic/gluconic acid distribution and abundance increased with distance from the organic PDA nutrient inoculation port, while the other organic acids were produced near and around the nutrient inoculation port (Figs. 2, S2, and S4). We also observed an increase in relative concentration of fungal organic acids in 30 days of fungal growth compared to 7 days of growth within the mineral doped micromodels (Fig. 2).

**Fig. 2.**
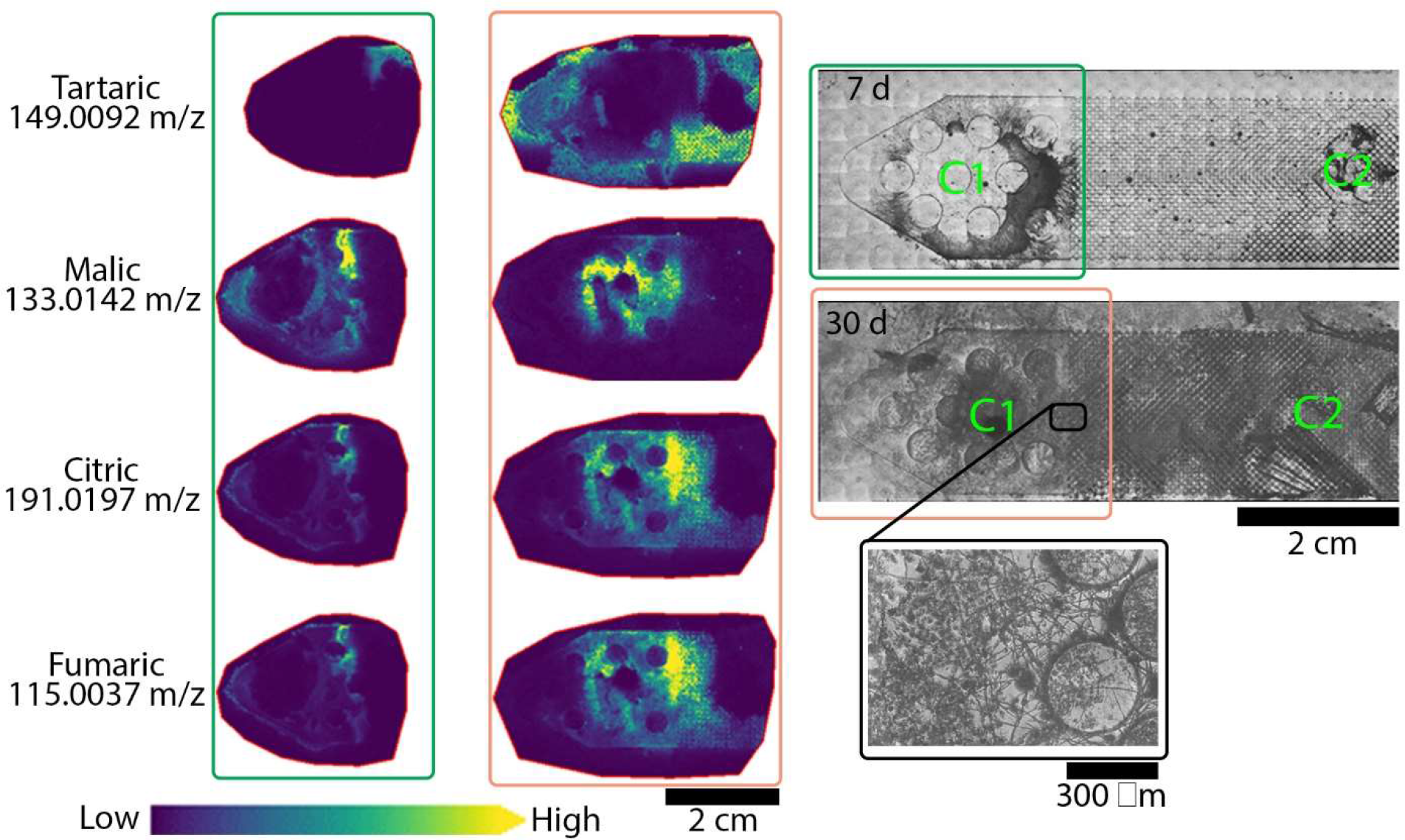
Heterogeneous distribution of fungal organic acids on mineral doped micromodel surfaces, as visualized with mass spectrometry. MALDI-MSI shows different organic acid production after 7 days (green box, left) and 30 days (orange box, center) of fungal growth on micromodels surfaces. The corresponding optical images show fungal growth on OG 603 mineral doped micromodels after 7 days (green box, top right) and 30 days (red box, center right). Fungi was inoculated along with PDA plugs at C1. C2 is the second carbon rich nutrient hotspot with only PDA plugs. The presence of minerals on the polymer surface decreases visibility of fungal hyphae, but hyphal growth is demonstrated across mineral grains (bottom right).

### Fungal organic acid uptake K into hyphae from mineral interfaces

To determine and visualize mechanisms of K uptake and transport by the fungal mycelia, the hyphal networks were removed from the surface of the soil micromodels through a polydimethylsiloxane (PDMS)-based embedding technique. This allowed us to analyze the K within fungal hyphae without interference from any mineral associated K on the micromodel surface using multi-energy micro-X-ray fluorescence imaging (μ-XRF), around the potassium K-edge, combined with potassium X-ray Absorption Near Edge Structure (XANES) spectroscopy. Here, we identified two distinct K chemistries specifically within fungal hyphae networks grown in the presence of soil minerals: organic acid-bound K and an unidentified form of K – ‘*organic K’* (Fig. 3). The unidentified *organic K* is present in all fungal hyphal biomass examined to date using XANES spectroscopy(13), with lesser contributions of organic acid-bound K. The unidentified *organic K* was observed in fungal hyphae grown on both mineral doped and mineral free micromodels. However, we observed K-citrate within hotspots along individual hypha and K-tartrate as mixtures with the unidentified *organic K* within the fungal hyphae from mineral doped micromodels alone (Fig. 3). The K-citrate and K-tartrate distribution in fungal hyphae is much lower than the unidentified *organic K*, which suggest fungi uptake and assimilate K into other forms such as *organic K* for storage in the hyphae. It is worth mentioning that identification of K citrate and K tartrate was made through spectral comparison of available synthetic standards. We observed that *Fusarium sp. DS 682* exuded several other organic acids such as malic acid and fumaric acid. These acid distributions were observed using MALDI MSI as well as GCMS, however due to the synthetic forms of these standard not being available, we were not able to confirm the contributions of other organic acids binding to K within the hyphae.

**Fig. 3.**
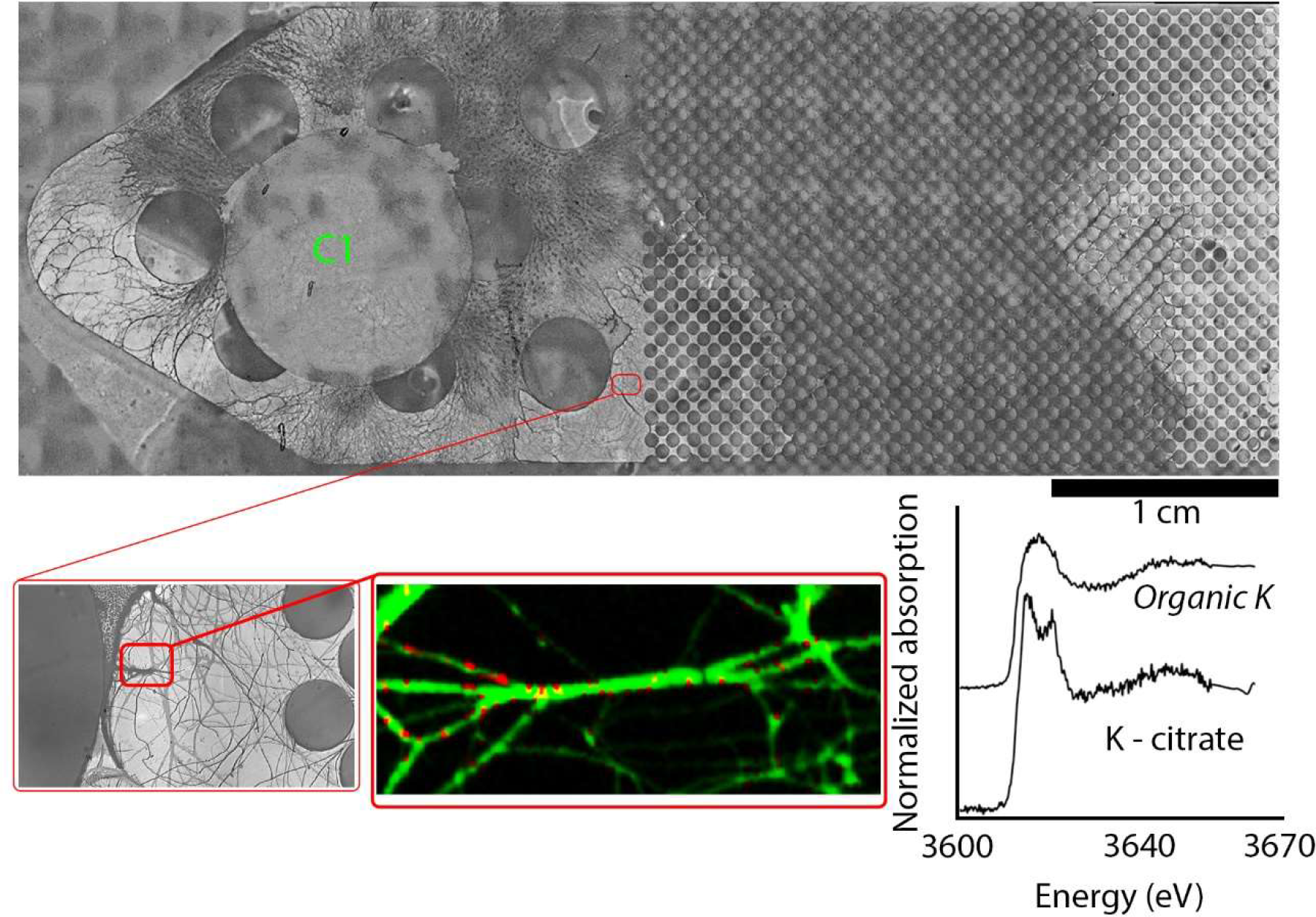
Fungi uptake K from mineral interfaces at hotspots along the hyphae. Fungal hyphae removed from mineral doped micromodel and embedded in PDMS (top image). C1 denotes MALDI MSI image around the inoculation/nutrient plug as shown in Fig.1. XANES spectroscopy from a region on micromodel (red box, bottom left) indicate the presence of two different forms of organic potassium; organic acid-bound K and an unidentified *organic K* used in XRF map fitting (Bottom, center). XANES spectra of *organic K* and organic-acid bound K.

### Fungi uptake K from minerals through organic acid chelation

The form of organic acid-bound K was analyzed by comparing XANES spectroscopy features of standards of organic acids (observed through analysis of fungal biomass growth on agar mineral surfaces; Fig. 1A) with the hyphal hotspots using linear combination fitting (LCF). The LCF analysis revealed that the organic acid-bound K hotspots contained different amounts of K-tartrate, K-citrate, and *organic K* (Fig. 4 and Table 1). Analysis of two positions on the hyphal biomass corresponding to the micromodel channel layout (Fig. 4A and B), using both μ- XRF maps and XANES spectroscopy, demonstrated a correlation between fungal organic acid production and location of organic PDA nutrient source. The XANES spectra obtained at several positions (A1-4 and B1-4) of the hyphal biomass was fitted with different standards (unidentified organic K, K-tartrate, K-citrate, and kaolinite; Figs. S5 and S6). Only organic K, and K-tartrate provided visually and statistically good fits (Fig. 4 and Table 1). These data show an increasing amount of K-tartrate away from the PDA nutrient source (area B Table 1). These results were further confirmed using nanospray desorption electrospray ionization mass spectrometry (nanoDESI-MS; Figs. S7 and S8). While accurate mass information from MALDI-MSI (Figs. 2, S2 and S3) and nanoDESI-MS techniques suggest the presences of these organic acid, nanoDESI was used with tandem mass spectrometry (MS/MS) to increase confidence in the annotation of these molecules. As shown in Fig. S8, malic and tartaric acid MS/MS data obtained from the fungal hyphae (measured directly from the microfluidic device) is highly comparable to data from authentic standards. This high degree of similarity between the experimental and authentic datasets (Pearson correlation coefficients of 0.98 and 0.94 for malic and tartaric acid datasets, respectively) strongly suggests that the discussed organic acid annotations are correct.

**Table 1.**
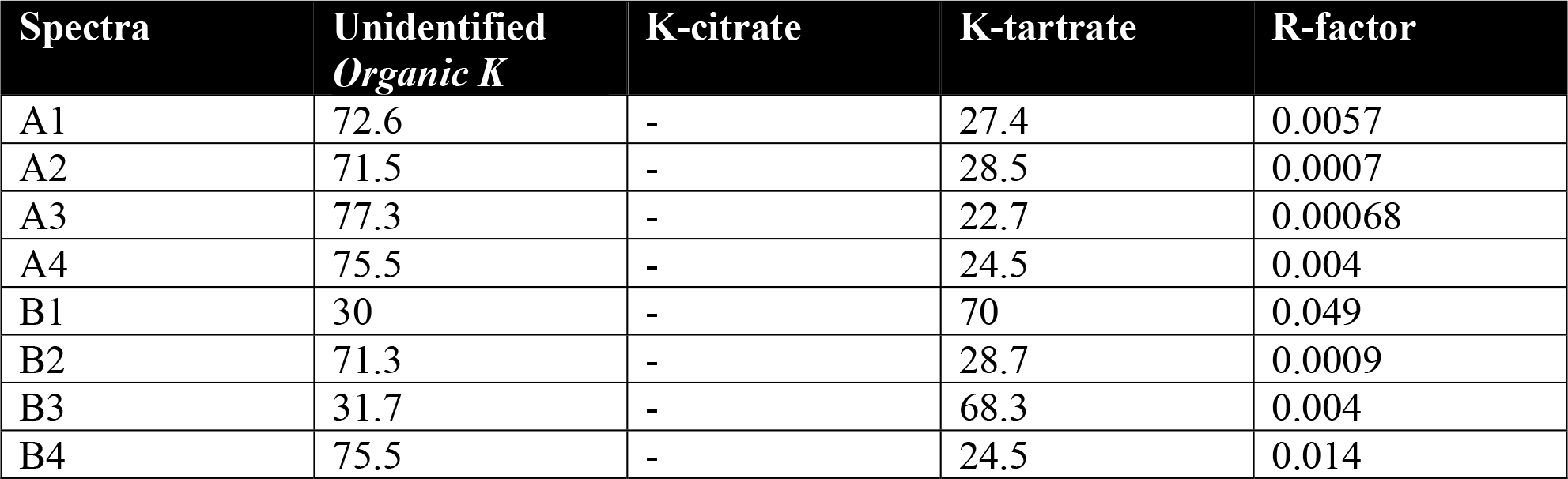
Spatial distribution of organic acid bound K changes with distance from carbon rich PDA. Linear combination fitting of the XANES spectra from Fig 4 (A1-4 and B1-4) shows amount (%) of *organic K*, K-citrate, and K-tartrate within fungal hyphae extracted from the micromodel surface.

**Fig. 4.**
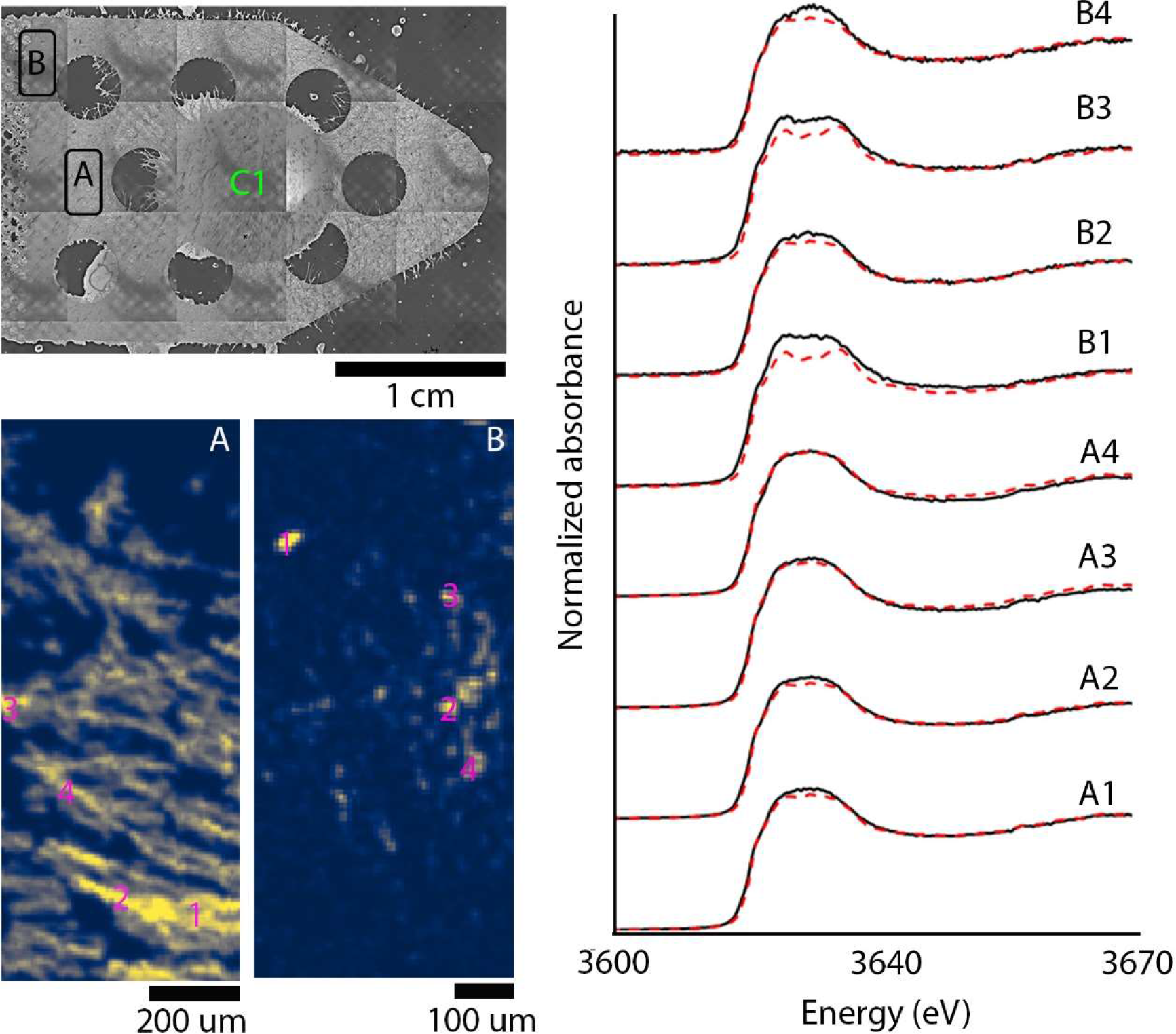
Uptake of K depends on the organic acid chelated as a function of distance from carbon hotspot. Two XRF maps show differences in K content within fungal hyphae from closer to the inoculation point (A) versus closer to the smaller pore space (B). Not only is there 20x less K counts in area B, but K also appears more as hotspots rather than along individual hyphae. Spectroscopy from region A is predominantly unidentified organic K while those from region B contain contributions of K-citrate and to a lesser extent K-tartrate. A fitting of the end-member spectra to the XRF maps of A and B show that only a small region within B has enough K-citrate that can be spatially mapped

## Discussion

The mechanisms involving microbially exuded small molecular weight acid dissolution and uptake of mineral derived nutrients is not well studied, partly due to the challenge of resolving such mechanisms in the complex soil landscape. However, these uptake mechanisms demonstrate how inorganic nutrients are cycled in terrestrial ecosystems as well as fate of certain nutrients in soil. In this work, we elucidate an indirect mineral weathering and nutrient uptake mechanism in the saprotrophic fungus, *Fusarium DS 682*, using a reduced complexity mineral doped soil micromodel platform. The MALDI-MSI compatible epoxy mineral doped soil micromodel system(15) provided us with a unique opportunity to spatially image fungal growth with and without soil minerals and correlate this with the spatial distribution of fungal organic acid exudates (Figs. 1B and 2). We observed lower relative concentrations of fungal organic acid in micromodels without minerals in comparison to micromodels with minerals, which suggest an increase in organic acid production in the presence of minerals (Figs. 2 and S2 and S3). The increase in fungal organic acid production in the presence of minerals is observed in both mineral doped micromodels and in fungal biomass from growth on agar plates (Fig. 1A). This could be attributed to the increased expression of organic acid transporter proteins in presence of minerals as observed in our previous work(13). In particular, the mineral doped micromodel platform facilitates greater resolution of how organic acid production changes with distance from a rich organic PDA source during fungal mineral weathering processes.

Furthermore, we observed differences in the distribution of the detected organic acids on the mineral doped micromodel surface that suggest these molecules perform different functions related to mineral weathering and nutrient uptake during fungal growth and nutrient foraging through soil minerals (Fig. 2). For example, citric acid, fumaric acid, and malic acid are part of the citric acid cycle and were abundant near the carbon rich PDA source. The production of these three organic acids perhaps occurs from degradation of dextrose in PDA via the citric acid cycle, in effort to release stored energy in the form of ATP, as well as produce NADH and amino acids precursors used in numerous biochemical reactions. On the other hand, tartaric acid, oxoadipic acid, and galactonic/gluconic acid (undistinguishable isomers) distribution was observed away from the PDA source, where nutrient limiting conditions predominated (Fig. S2). Given microorganisms, such as fungi, possess mechanisms for storing energy to survive in the event of limited environmental resources, and glycogen represents a primary energy storage form in fungi, we propose the stored glycogen in the fungal biomass is metabolized for biosynthesis of acids (e.g., tartaric acid) away from the rich PDA nutrient source. Therefore, as *Fusarium sp. DS 682* is growing away from the nutrient source, into a carbon limited environment utilizing a different metabolic pathway to generated chelating organic acids. Notably, tartaric acid biosynthesis does not produce ATP and is perhaps only synthesized as a means to extract K from mineral interfaces. A product of glycogen metabolism is glucose, and we observed hexose/glucose distribution colocalized with tartaric acid distribution (Fig. S9), which suggest glycogen was metabolized to create tartaric acid, oxoadipic acid, and galactonic/gluconic acid. It is known that oxoadipic acid and galactonic/gluconic acid are produced during biosynthesis of tartaric acid(16). We also observed several other intermediate products of tartaric acid biosynthesis pathway such as ascorbate, dehydroascorbate, and iodonate(16) colocalized with tartaric acid away from carbon rich PDA hotspot (Fig. S10). Combined, the production of such intermediate products and organic acid suggests that there is a switch in organic acid production away from a rich carbon hotspot such as the PDA.

Comparison of Pearson’s correlation coefficients between citric acid, tartaric acid, and hexose spatial distribution demonstrates a weak correlation between tartaric acid and hexose and a weaker correlation between citric acid and hexose at 7 days of fungal growth. However, the correlation between tartaric acid and hexose increases at 30 days of fungal growth, while the correlation between citric acid and hexose distribution remains weak (Table S1). These distinct difference in acid production demonstrates that fungal organic acid production in the presence of soil minerals is influenced by spatially distributed carbon rich hotspots. We observed that the carbon rich PDA plugs are used by fungi within 7-14 d of growth. However, fungal growth and organic acid production was observed for up to 30 d, with increase in tartaric acid production within the soil like pore spaces away from the PDA source (Fig 2 and S2). Therefore, the increased production of organic acids with time is perhaps for increasing nutrient extraction from mineral interfaces in a low carbon environment.

We previously demonstrated increased transport of K through fungal hyphae from the natural kaolinite mineral interfaces of mineral doped micromodels (13). Here, we analyzed K bonding environment within fungal hyphae after growth with and without minerals using XRF maps and XANES spectroscopy. There are several recent spectroscopic investigations demonstrating K chemistry in soils (17, 18) and leaf litter (18), however spectroscopic studies of K speciation within biological systems are limited. Therefore, K -spectra observed here had to be correlated with spectra of available standards. We observed two different K chemistries, *organic K* and organic acid bound K (specifically, K-citrate and K tartrate) (Fig. 3 and 4), where K citrate were present in distinct hotspots along fungal mycelia. The *organic K* is likely a mixed phase where there are contributions from K-citrate, K tartrate, other organic acid bound K and other organic compound bound K. We observed the organic K spectral features in the hyphae in the control without minerals. This suggest that the *organic K* spectra is complex and has contributions from several different K-chemistries. We are able identify K tartrate and K citrate through spectral comparison of synthetic standards, however we do not rule out contributions from other K bonded compounds to the *organic K*. While the organic acids demonstrate a distance dependent exudation from the carbon rich nutrient source as observed by MALDI/MSI and NanoDESI, the distribution of the K chelated acids within the hyphae were uniform (Fig. 3 and 4, Table 1). XRF imaging and XANES spectra of fungal biomass separated from the micromodels demonstrated K complexed with tartaric acid existed in hyphae growing in areas where no tartaric acid exudation was observed using MALDI MSI and nanaoDESI. Moreover, citric acid complexed K was found only in hotspots within the hyphae. These results suggest that the K-complexed with organic acids were being transported through the fungal hyphae and perhaps being stored for future use (Fig. 4 and S6). Even though there is no distance dependance on uptake, we observe a distance dependence exudation of organic acids that suggest fungal organic acid production was for indirect weathering of minerals for uptake of mineral derived nutrients. Moreover, the K-citrate and K-tartrate distribution in fungal hyphae was much lower than *organic K* (Fig. 3 and 4, Table 1), which suggest fungi uptake and assimilate K into a different form, such as for storage in the hyphae. This stored K form here corresponds to *organic K* that we observe in fungal hyphae growth in both mineral-doped and mineral-free micromodels. The increased spatial distribution of citric acid and tartaric acid on mineral-doped micromodel surfaces, as well as the presence of K-citrate and K-tartrate in hyphal biomass of fungi grown in mineral-doped micromodels, combined that suggest fungi chelate and uptake K from mineral using organic acids through an indirect mineral weathering and uptake mechanism.

The role of fungal produced organic acids has been suggested in the uptake of mineral derived nutrients in the past(19). In this study, we visualize and demonstrate the organic acid uptake of mineral drive K by fungal hyphae with unprecedented spatial and molecular resolution. Further, we show that the production of the type of fungal organic acids is a function of distance from rich carbon hotspot and the organic acid uptake of K correlate to this spatial distribution of exuded organic acids. Biotic mineral degradation is one of the biogeochemical processes that regulate the flux of mineral derived inorganic nutrient pool. Mineral derived inorganic nutrients like K cycles through soil organic matter and provide much required nourishments to maintain life in terrestrial ecosystems, especially in nutrient impoverished soils. Such inorganic nutrient pools also influence the composition of carbon hotspots in soils and microbial community development resulting from proximity to such hotspots. This study provides a novel mechanistic insight into formation of inorganic-organic nutrient complexes in fungal biomass during an indirect mineral weathering process, that can be used as a framework toward unraveling biotic driven mineral weathering mechanisms.

## Methods

### Fungal strain and growth media

The soil fungus *Fusarium sp. DS682* (14) was cultured as previously described (13). Briefly, *Fusarium sp. DS682* was cultured on potato dextrose agar (PDA) plates by inoculating 100 μl of spores from frozen stocks. The inoculated PDA plates were incubated at 28° C for 15 days before removing fungal biomass for polydimethylsiloxane (PDMS)-glass and PDMS-epoxy micromodel growth.

### Microfluidics device fabrication

We utilized two different mineral doped micromodel devices: a PDMS-glass and PDMS-epoxy based-devices. Both devices were created from a silicon master as described before (13). Briefly, the microfluidics devices for MALDI-MSI were constructed using a combination of the Bosch process (20) and soft lithography techniques (21). A positive silicon (Si) master of the microfluidic channel that was previously described (22) was constructed using the Bosch process and functionalized using chlorotrimethylsilane (CTMS; Sigma Aldrich, St. Louis, MO) by exposure in a desiccator under vacuum overnight. Negative replicas of the Si master were produced from PDMS (Sylgard 184, Dow Corning, Midland, MI) with a prepolymer-to-curing agent ratio of 10:1. After extensive mixing of the prepolymer and curing agent, the mixture was poured on the Si master of the microchannel and placed in a vacuum desiccator for 1 h to eliminate all air bubbles. It was then thermally cured in an oven for 3 h at 70° C. After cooling, the negative PDMS mold was gently peeled off the Si substrate. The negative PDMS mold was then cleaned extensively with ethanol, isopropanol, and acetone sequentially, dried and treated in oxygen plasma for 30 s in a plasma cleaner (PX250, Nordson March, Concord, CA). After this surface treatment, the PDMS negative mold was placed in a CTMS environment in a desiccator under vacuum overnight.

In order to produce the final positive replica of the microchannel, glass coverslips were functionalized using aminopropyltrimethoxysilane (APTMS; Sigma Aldrich, St. Louis, MO). Devices were placed in APTMS environment in a vacuum desiccator overnight. A layer of OG603 epoxy (Epoxy Technology Inc., Billerica, MA) was spin coated using a WS-400B-6NPP/LITE spin coater (Laurell Technologies Corp., North Wales, PA) to the functionalized glass coverslip at a spin speed of 500 rpm for 10 s. Then, the PDMS negative channel was filled with OG603 epoxy and placed under vacuum in a desiccator to ensure the air bubbles are eliminated and the epoxy filled the entire microchannel space. The microchannel surface with epoxy was placed against the epoxy coated glass coverslip and pressed to remove any excess epoxy. This set up was then cured under UV light (Melody Sussie UV gel nail polish dryer) overnight to solidify. Once the epoxy material had solidified, the negative PDMS replica was peeled off, leaving the epoxy microchannel positive with a glass coverslip backing. This process produced epoxy films that are 5-7 μm thick and requires a glass coverslip backing for stability. These epoxy microchannels are compatible with MA(LDI) and optical microscopy techniques (15).

### Microfluidic device assembly and fungi inoculation

The epoxy microfluidic channels were plasma cleaned and reversibly bonded to a 5 mm PDMS layer with ports at two ends for inoculation of Fungal biomass For reversible bonding of PDMS to epoxy, a 5 mm PDMS layer was created by pouring 40 g of prepolymer-to-curing agent ratio of 20:1 in a 150 × 15 mm petridish (VWR, Radnor, PA) and cured at 70° C for 30 min. Creating inoculation port covers for the devices followed the same 20:1 ratio PDMS recipe, and 20 mL of this mixture were poured into a 150 mm diameter petri dish and cured at 70° C for 30 min. The cured polymer was cut into 6 mm × 6 mm squares and plasma cleaned (PX250, Nordson March, Concord, CA) before using to seal inoculation ports of the micromodels.

### Gas chromatography mass spectrometry (GC-MS) sample preparation and analysis

Fungal samples for metabolomics analysis were extracted from fungal biomass grown on PDA agar. To expose the fungus to minerals, 100 μl of 50% (w/v) natural kaolinite solution in water were autoclaved and spread over the agar surface using a disposable L shaped spreader (Thermo Fisher Scientific, Waltham, MA). The solution was dried for 1 h to create a thin mineral film on the agar surface. Three plates each with (+ mineral) and without mineral (- mineral) condition were inoculated with fungal biomass from 14 d growth on PDA using a 1.5 mm holepunch. The plates were incubated at 28° C for 15 d before extracting hyphae (500 mg) from each plate. The metabolite samples were extracted using the MPLEx protocol as previously described (23). A solution of 2:1 CH3Cl:CH3OH was added at 5 times the volume of the cell lysate and vortexed for 30 s and kept on ice for 5 min and vortexed again for 30 s. The solution was centrifuged at 10000 x g for 10 min at 4° C and the upper aqueous layer was collected for metabolite analysis in a 2 ml flat bottle autosampler vial. The metabolite part was then dried overnight in a speedvac and stored in – 80° C for further analysis.

Dried extracts were chemically derivatized using a modified version of the protocol used to create FiehnLib (24). Briefly, dried metabolite extracts were dried again to remove any residual water from being stored at −80 °C. To protect carbonyl groups and reduce the number of tautomeric isomers, 20 μl of methoxyamine (Sigma Aldrich) in pyridine (Sigma Aldrich) (30 mg ml^−1^) was added to each sample, followed by vortexing for 30 s and incubation at 37 °C with vigorous shaking (1,000 rpm) for 90 min. The sample vials were then inverted once to capture any condensation of solvent at the cap surface, followed by a brief centrifugation at 1,000 × *g* for 1 min. To derivatize hydroxyl and amine groups to trimethylsilylated forms, 80 μl of *N*-methyl-*N*-(trimethylsilyl)trifluoroacetamide (Sigma Aldrich) with 1% trimethylchlorosilane (Sigma Aldrich) were then added to each vial, followed by vortexing for 10 s and incubation at 37 °C with shaking (1,000 rpm for 30 min. Again, the sample vials were inverted once, followed by centrifugation at 1,000 × *g* for 5 min. The samples were allowed to cool to room temperature and analyzed the same day.

An Agilent GC 7890A coupled with a single quadrupole MSD 5975C (Agilent Technologies) was used and the samples were analyzed in randomized fashion. An HP-5MS column (30m × 0.25 mm × 0.25 μm; Agilent Technologies) was used for untargeted metabolomics analyses. The sample injection mode was splitless and 1 μl of each sample was injected. The injection port temperature was held at 250 °C throughout the analysis. The GC oven was held at 60 °C for 1 min after injection and the temperature was then increased to 325 °C by 10 °C min^−1^, followed by a 5 min hold at 325 °C (25). The helium gas flow rates for each experiment were determined by the Agilent Retention Time Locking function based on analysis of deuterated myristic acid and were in the range of 0.45–0.5 ml min^−1^. Data were collected over the mass range 50–550 *m/z*. A mixture of fatty acid methyl esters (C8–C28) was analyzed once per day together with the samples for retention index alignment purposes during subsequent data analysis.

GC-MS raw data file processing was done using Metabolite Detector software and metabolites were identified by matching experimental spectra and retention indices to an augmented version of FiehnLib (24). All identifications were manually validated to reduce deconvolution errors and to eliminate false identifications. The NIST 14 GC–MS library was also used to cross-validate the spectral matching scores obtained using the Agilent library and to provide identifications of unmatched metabolites.

### MALDI-MSI sample preparation and analysis

Microfluidic devices were sprayed with 7 mg/mL of N-(1-naphthyl) ethylenediamine dihydrochloride (NEDC; Sigma Aldrich) dissolved in 70% MeOH using a TM-Sprayer (HTX Technologies). Spraying conditions were 8 passes at 1,200 μL/min, nozzle temperature 75°C, a spray spacing of 3 mm, and a spray velocity of 1300 mm/min. MS imaging analysis was performed on a 15 Tesla MALDI-FTICR-MS (Bruker Daltonics) equipped with SmartBeam II laser source (355 nm, 2 kHz) using 200 shots/pixel with a frequency of 2 kHz and a 150 μm step size. FTICR-MS was operated in negative mode to collect ions with m/z 92-700, using a 209 ms transient, that translated to a mass resolution of R ∼ 60,000 at 400 m/z. Data was acquired using FlexImaging (v 4.1, Bruker Daltonics), and image processing, segmentation, co-localization analysis, overlaying with optical images and visualization were performed using SCiLS Lab (Bruker Daltonics).

### Optical microscopy

Fungal growth in the micromodels were imaged using a Nikon eclipse TE2000-E epifluorescence microscope (Nikon, Melville, NY) using a 4× air objective. The micromodels were imaged by acquiring a mosaic image of the sample surface. The separate tiles were then compiled and imported into an image processing program (Fiji) (26) as an image stack. The rolling ball BASIC plugin (27) was used to remove vignetting in individual tiles, then image stitching was performed to recombine the individual image tiles into a mosaic image from multiple image tiles. Mosaic brightfield images were obtained and stitched across the length and width of the microchannel to capture fungal growth in micromodels.

### SEM and EDX analysis of epoxy substrates

Sections were cut out from epoxy micromodels for SEM analysis to fit into the SEM sample holder. All micromodel sections were mounted into aluminum stubs. To reduce sample charging, the sections were coated with∼15 nm carbon using a thermal evaporation method (108C Auto Carbon Coater, Ted Pella, Inc.) and were secured with Cu sticky tape on the SEM stage. SEM analyses were conducted with a FEI Helios NanoLab 600i field emission electron microscope. Images were collected with Everhart-Thornley secondary electron detector (ETD) in a field free mode and at acceleration voltage of 3 to 5 kV, current of 0.086 to 0.17 nA and∼ 4 mm working distance.

The chemical composition of mineral particles embedded in the epoxy and elemental distribution maps, which illustrate distribution of mineral grains, were collected with an energy dispersive X-ray detector (EDX). The SEM was equipped with an EDX X-Max 80mm^2^ Silicon Drift Detector (Oxford Instruments, Abington, UK) capable of detecting lighter elements. EDX analyses were performed at 3 kV to visualize C-elemental distribution and at 10 kV to collect chemical data on K, Ca, Mg, Si, and Al. Oxford INCA software was used to collect compositional maps (acquisition time > 300 s) and point spectrum analyses (acquisition time∼80 s).

### Synchrotron XRF imaging and XANES spectroscopy

Fungal hyphae removed from micromodel surfaces were embedded in PDMS in order to map K associated with hyphae (and not minerals). Potassium K-edge μ-XRF maps and XANES spectroscopy were collected at beamline 14-3 at the Stanford Synchrotron Radiation Lightsource (SSRL), SLAC National Laboratory. The double Si crystal (111) monochromator was calibrated using a natural kaolinite powder (Sigma Aldrich, S. Louis, MO) to 3618.73 eV (13). An X-ray spot size of 1 micron was focused by Sigray achromatic paraboidal lenses. Samples were loaded onto a cassette and placed within a multi-sample wheel, and subsequently measured at room temperature in a helium purged atmosphere. Multiple energy μ-XRF mapping at various energies around the K K-edge, combined with XANES spectroscopy, can spatially resolve different K chemistries across the mapped region. The energies for mapping were 3615.70, 3617.25, 3618.50, and 3621.80 eV, which correspond to features observed in the XANES spectra of both standard materials and previous experiments. A principal component analysis and simplex volume maximization on the multi-energy maps were used to provide information on where different K chemistries are located across the mapped region. These locations were selected for XANES spectroscopy in order to identify the chemistry present. Standards for XANES spectroscopy (tri-potassium citrate monohydrate and potassium tartarate) were prepared as finely milled powders and spread as a thin layer onto sulfur free tape, with excess powder tapped off. Spectra of the standards were collected using an unfocussed beam (∼1mm × 2 mm) with at least 3 repeat scans.

Data were processed using SMAK for μ-XRF maps (28) and SIXpack (29) and Athena for XANES spectroscopy (30). XANES spectra were processed by averaging repeat scans, background subtracted and normalized using standard protocols (13). A linear combination fitting (LCF) of standard XANES spectra to the experimental XANES was used to model the relative proportions of various K chemistries within a single spot on a sample. To determine the spatial relationships of the chemistries identified to be present by LCF, XANES standards were fit to the multiple energy μ-XRF maps. Multi-energy maps are imported to a single file and background noise is reduced at each energy by applying a Gaussian distribution function of a 5 pixel area known as blurring. Speciation maps are generated by applying a least squares fitting of standard XANES spectra to the blurred multi-energy maps.

### nanoDESI-MS analysis

Related nanospray desorption electrospray ionization (nano-DESI) MS sampling apparatuses have been described in detail elsewhere (8, 31, 32). Two silica capillaries (150 μm O.D., 50 μm I.D.; Polymicro Technologies, Phoenix, USA) were aligned to meet at a ∼90° angle. Solvent (7/3 methanol/deionized water v/v) was flowed through the ‘primary’ capillary at 1 μL/min via a syringe pump, whereas the nano-DESI capillary assembly was positioned such that the tip of the secondary capillary was positioned ∼1 mm from the inlet of a Thermo LTQ Orbitrap Velos high resolution mass spectrometer (operated in negative ion mode). All studies were conducted at a mass resolution setting of 100k at *m/z* 400. A potential of -3.5 kV was applied to the solvent via the syringe needle. The junction between the primary and secondary capillaries was brought sufficiently close to the sample surface such that solvent bridge was formed, which was then scanned across the sample surface at 30 μm/s for spatial analyses. The MS inlet was maintained at 300°C for all analyses. For MS1 studies, the MS was operated with a scan range of *m/z* 100 – 600, a maximum ion injection time of 200 ms, and an automatic gain control (AGC) target of 1×10^6^. For MS/MS studies, the MS was operated with an MS1 isolation width of 0.1 amu, a MS2 scan range of *m/z* 50 – 150, a maximum ion injection time of 1000 ms, and an automatic gain control (AGC) target of 5×10^5^. Collision energies of 21 and 24 (arbitrary units) were used for malic and tartaric acid analyses, respectively.

## Acknowledgements

This research was performed on a project award (<u>10.46936/intm.proj.2021.60094/60001433</u>) from the Environmental Molecular Sciences Laboratory, a DOE Office of Science User Facility sponsored by the Biological and Environmental Research program under Contract No. DE-AC05-76RL01830. Use of the Stanford Synchrotron Radiation Lightsource (SSRL), SLAC National Accelerator Laboratory, is supported by the DOE, Office of Science, Office of Basic Energy Sciences under Contract No. DE-AC02-76SF00515. The SSRL Structural Molecular Biology Program is supported by the DOE-BER, and by the National Institutes of Health (NIH), National Institute of General Medical Sciences (NIGMS, P30GM133894). The contents of this publication are solely the responsibility of the authors and do not necessarily represent the official views of NIGMS or NIH. A portion of this research was supported by the U.S. Department of Energy (DOE) Office of Biological and Environmental Research (BER) and is a contribution of the Scientific Focus Area “Phenotypic response of the soil microbiome to environmental perturbations.” Pacific Northwest National Laboratory (PNNL) is operated for the DOE by Battelle Memorial Institute under Contract DE-AC05-76RLO1830.

## Contributions

AB and CRA designed the experiments and wrote the paper. AB performed all the experiments. OQ performed SEM analysis. JR performed the XANES and XRF analysis. DV performed the MALDI-MSI analysis. SPC performed the GCMS analysis. GWV performed the nanoDESI-MS analysis. MJT performed optical microscopy image analysis. All authors edited the manuscript, with AB and CRA providing the main edits.

## Ethics declaration

The authors declare no competing interests

## Notes

### Competing Interest Statement

The authors have declared no competing interest.

