## Supplementary figures and tables for "Fungal organic acid uptake of mineral derived K is dependent on distance from carbon hotspot"

**Table of Contents:**

**Supplementary figures**

**Fig. S1.** Potassium mineralogy is emulated within mineral doped micromodels ..……..……….3

**Fig. S2.** Organic acid distribution changes away from a rich carbon hotspot…………………..……4

**Fig. S3**. Fungal organic acid production decreased on mineral free micromodels……………..….5

**Fig. S4.** Fungal organic acid distribution and abundance changes with distance from the carbon rich nutrient source…………………………………………………………………….……………6

**Fig. S5**. XANES spectra of standards used for comparison of peak energies observed in fungal hyphae ……………………………………………………………………………….….…....7

**Fig. S6**. *Organic K* is the predominant form of K in fungal hyphae………………….…...…………8

**Fig. S7.** Linear nanoDESI-MS line scans for measured relative abundances of organic acids…...9

**Fig. S8.** Tandem mass spectrometry (MS/MS) comparison data for malic and citric acids measured directly from fungal hyphae (microfluidic device) and authentic standards…………10

**Fig. S9**. Tartaric acid biosynthesis occurs through degradation of stored energy in fungi……...11

**Fig. S10**. Tartaric acid biosynthesis occurs away from carbon rich nutrient hotspot…………...12

**Supplementary Tables**

**Table S1**. Pearson's correlation coefficients from comparing distribution of hexose, citric acid, and tartaric acid from different stages of fungal growth on mineral doped micromodels………13

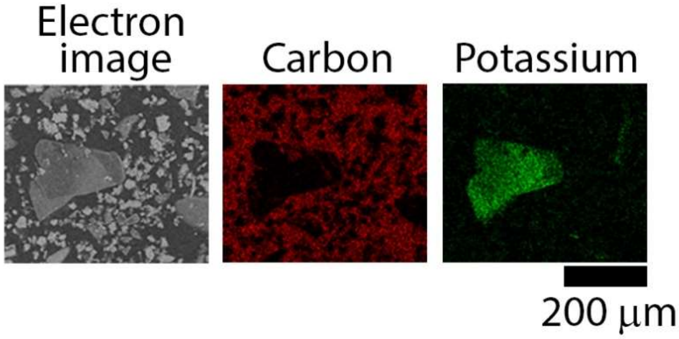


Fig. S1. **Potassium mineralogy is emulated within mineral doped micromodels.** Scanning electron microscopy and energy dispersive X-ray analysis demonstrates presence of K in mineral grains embedded on the micromodel surface.

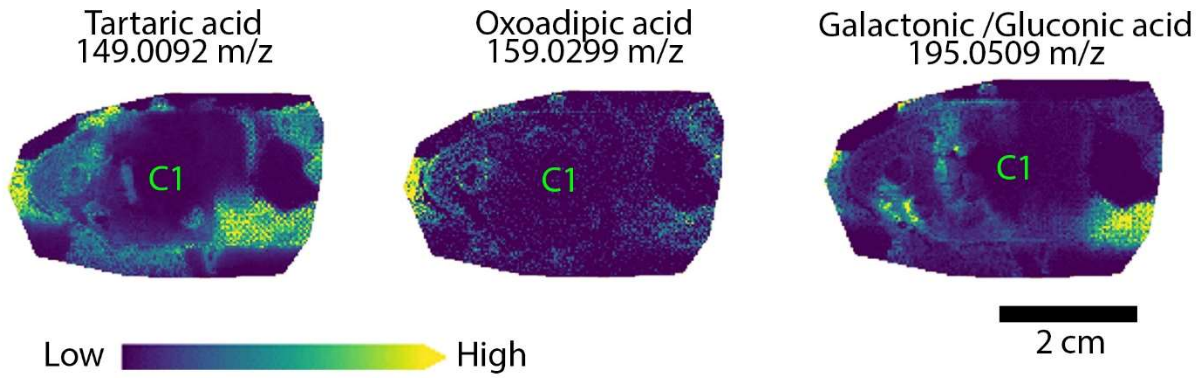


**Fig. S2. Organic acid distribution changes away from a rich carbon hotspot**. MALDI-MSI data shows distribution of tartaric acid, oxoadipic acid and galactonic/gluconic acid are similar and away from the carbon rich PDA source. C1 denotes MALDI MSI image around the inoculation/nutrient plug as shown in Fig.1.

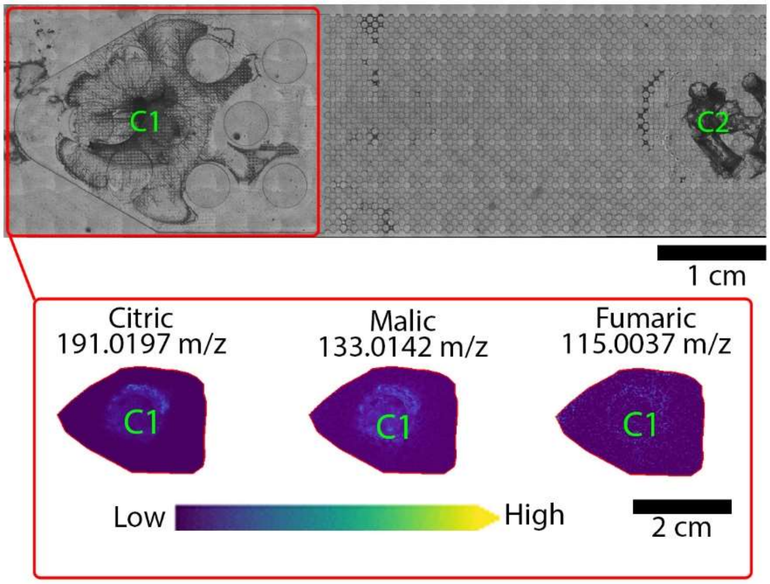


**Fig. S3. Fungal organic acid production decreased on mineral free micromodels**. Fungal hyphal growth on OG 603 micromodels without minerals (top image). MALDI-MS images demonstrate organic acid distribution on the micromodel surface after fungal growth (bottom panel of images). C1 denotes MALDI MSI image around the inoculation/nutrient plug as shown in Fig.1.

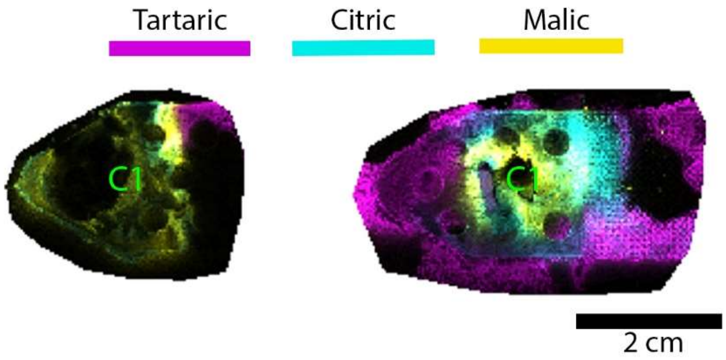


**Fig. S4. Fungal organic acid distribution and abundance changes with distance from the carbon rich nutrient source**. Fungal organic acid distribution on mineral doped micromodels after 7 days (left image) and 30 days (right image) of fungal growth. The tartaric acid, citric acid, and malic acid distribution shows tartaric acid production occurs away from the carbon rich PDA source. The spatial distribution of organic acids, such as citric acid and tartaric acid that is used to chelate K do not overlap. C1 denotes MALDI MSI image around the inoculation/nutrient plug as shown in Fig.1.

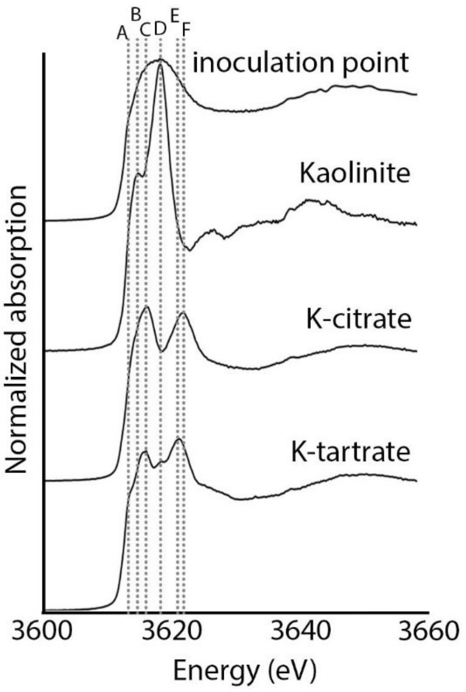


**Fig. S5. XANES spectra of standards used for comparison of peak energies observed in fungal hyphae.** XANES spectra was obtained at the potassium K-edge of K salts of organic acids (K-citrate and K-tartrate), kaolinite mineral, inoculation point (*organic K).* Grey dotted bars correspond to energies of unique, identifying features of the spectra; A – pre-edge feature on the rising edge of potassium tartarate at 3614 eV, B – pre-edge feature of natural kaolinite powder at 3615.4 eV, C – edge position of tripotassium citrate and edge feature of K-tartarate at 3616.5 eV, D – edge position of inoculation point and kaolinite, small edge feature of potassium tartarate at 3618.8 eV and E –edge feature of tripotassium citrate at 3622.1 eV and F – edge feature of tripotassium citrate at 3622.6 eV.

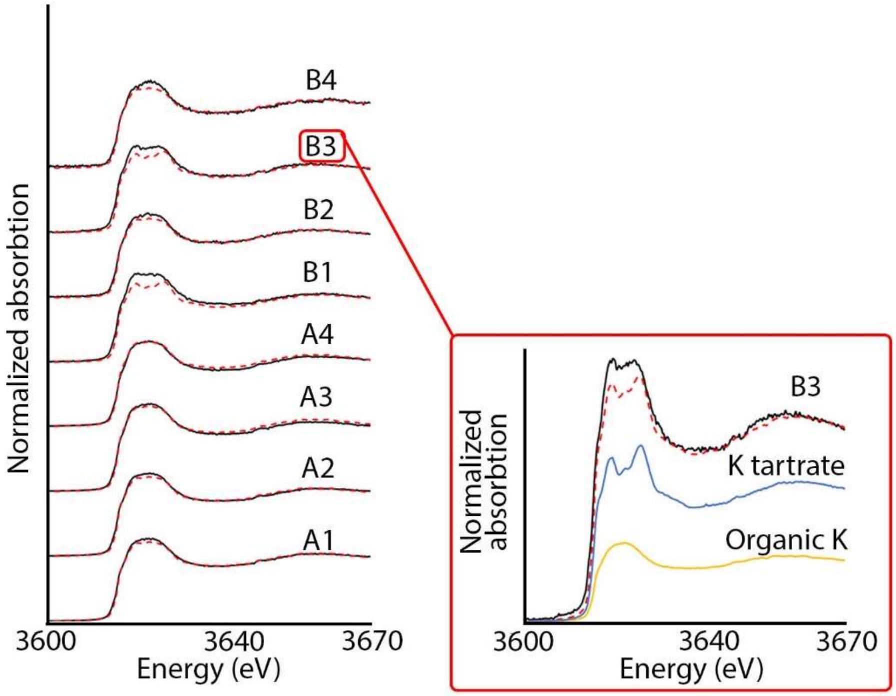


**Fig. S6. *Organic K* is the predominant form of K in fungal hyphae**. Linear combination fitting of XANES spectra from fungal hyphae using standard K salts and organic K shows amount of *organic K* and organic acid bound K within fungal hyphae (Table S1). Here, spectra 1, 2, 3, and 4 within the red box represents test B3 spectra with LCF overlay (red dotted spectra), K-tartrate and *organic K* respectively.

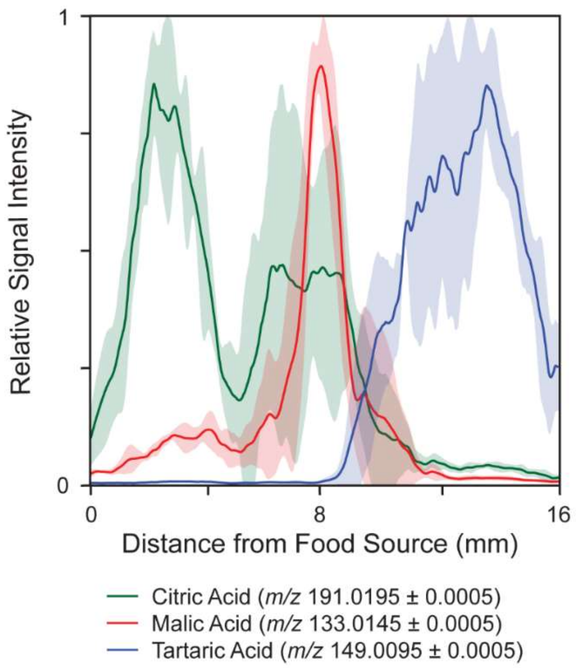


**Fig. S7. Linear nanoDESI-MS line scans for measured relative abundances of organic acids.** Shaded areas represent ± one standard deviation of triplicate line scans (0 mm represents edge of the food source). Data for each organic acid is normalized to the maximum signal intensity.

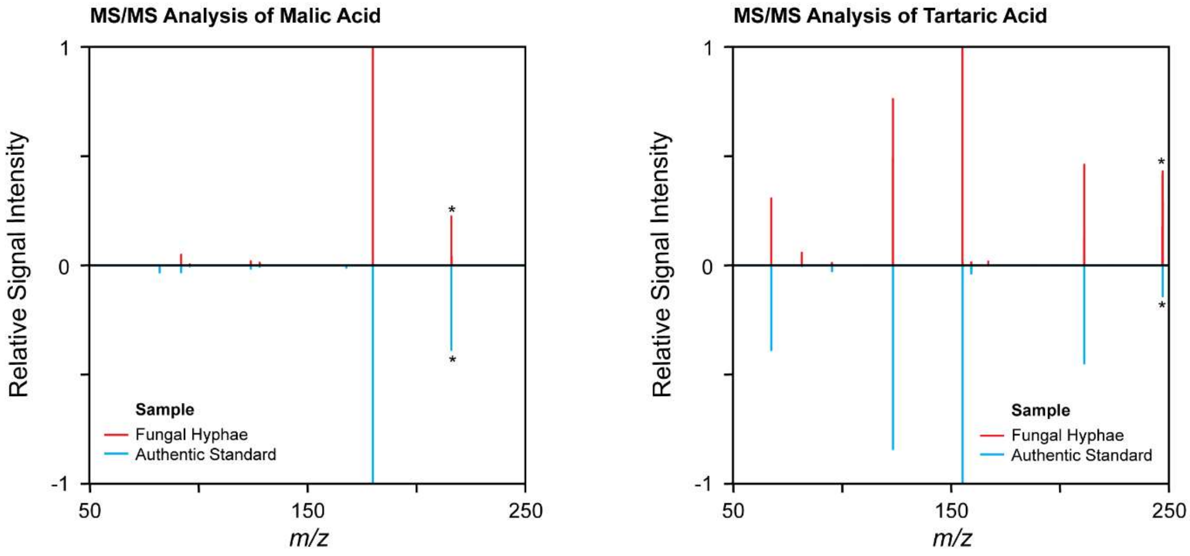


**Fig. S8. Tandem mass spectrometry (MS/MS) comparison data for malic and citric acids measured directly from fungal hyphae (microfluidic device) and authentic standards.** MS/MS spectra are an average of ≥15 scans. Pearson correlation coefficients as presented in the manuscript are for MS/MS features >4% relative intensity. Asterisks indicates MS1 masses for fragmentation.

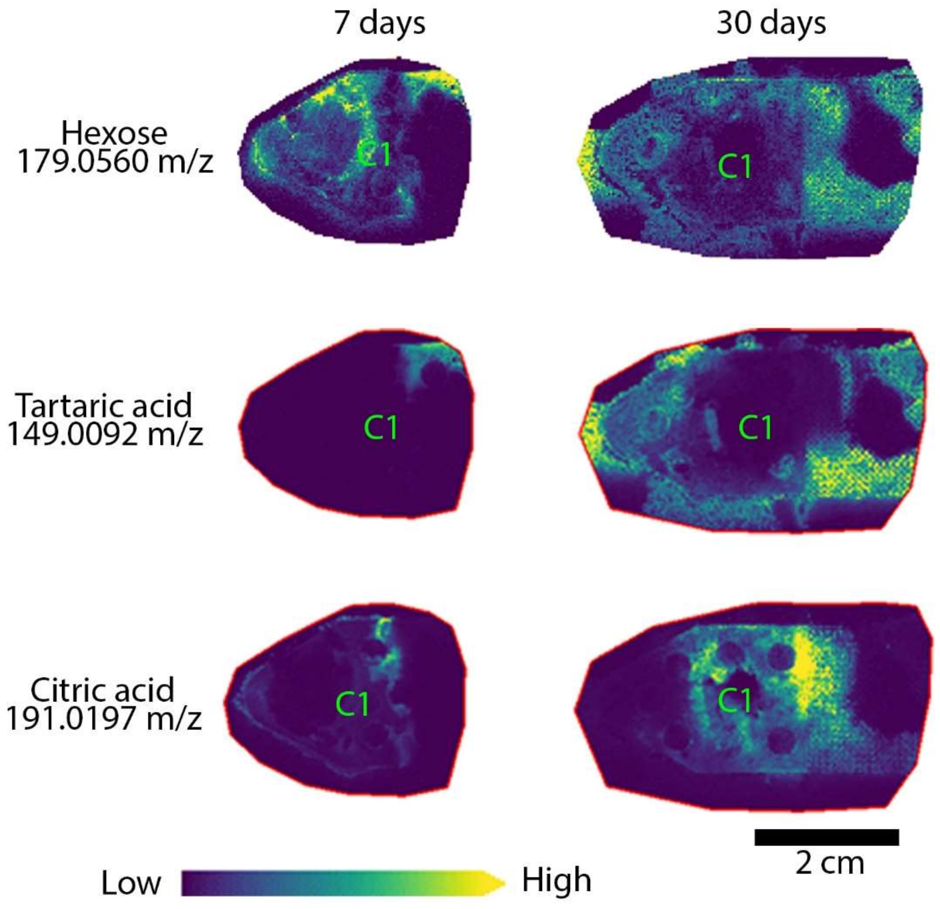


**Fig. S9. Tartaric acid biosynthesis occurs through degradation of stored energy in fungi.** MALDI-MSI demonstrates colocalization of hexose/glucose and tartaric acid distribution on the mineral doped micromodel after 30 days of fungal growth (see Pearson's correlation coefficient values in Table S1). The hexose distribution at 7 days of fungal growth is more similar to citric acid which is observed near to carbon rich PDA. The hexose/glucose production at 30 days perhaps occurs from degradation of stored glycogen in fungal hyphae. C1 denotes MALDI MSI image around the inoculation/nutrient plug as shown in Fig.1.

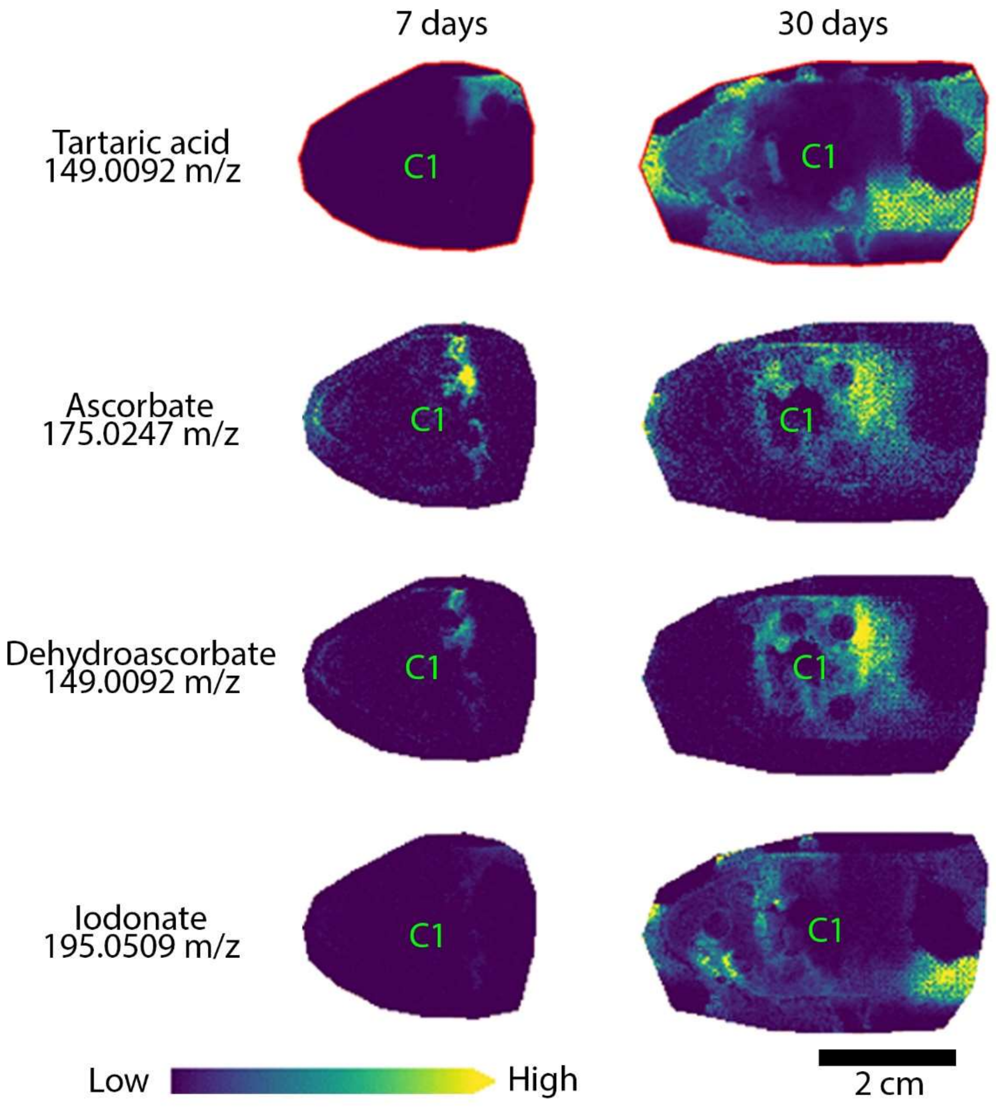


**Fig. S10. Tartaric acid biosynthesis occurs away from carbon rich nutrient hotspot**. MALDI-MSI shows distribution of intermediate products from the tartaric acid biosynthesis. The distribution of such intermediate products is colocalized with tartaric acid distribution on the mineral doped micromodel after fungal growth. C1 denotes MALDI MSI image around the inoculation/nutrient plug as shown in Fig.1.

**Table S1. Pearson's correlation coefficients from comparing distribution of hexose, citric acid, and tartaric acid from different stages of fungal growth on mineral doped micromodels.**

| Micromodel conditions | Pearson's correlation coefficient |
|---|---|
| 7 days citric vs 7 days hexose | 0.02 |
| 30 days citric vs 30 days hexose | 0.05 |
| 7 days tartaric vs 7 days hexose | 0.22 |
| 30 days tartaric vs 30 days hexose | 0.50 |
